# Conservation and divergence in modules of the transcriptional programs of human and mouse immune systems

**DOI:** 10.1101/286211

**Authors:** Tal Shay, Vladimir Jojic, Or Zuk, Christophe Benoist, Daphne Koller, Aviv Regev, ImmGen Consortium, Paul Monach, Susan A. Shinton, Richard R. Hardy, Radu Jianu, David Laidlaw, Jim Collins, Roi Gazit, Brian S. Garrison, Derrick J. Rossi, Kavitha Narayan, Katelyn Sylvia, Joonsoo Kang, Anne Fletcher, Kutlu Elpek, Angelique Bel-lemare-Pelletier, Deepali Malhotra, Shannon Turley, Adam J. Best, Jamie Knell, Ananda Goldrath, Vladimir Jojic, Daphne Koller, Tal Shay, Aviv Regev, Nadia Cohen, Patrick Brennan, Michael Brenner, Taras Kreslavsky, Natalie A. Bezman, Joseph C. Sun, Charlie C. Kim, Lewis L. Lanier, Jennifer Miller, Brian Brown, Miriam Merad, Emmanuel L. Gautier, Claudia Jakubzick, Gwendalyn J. Randolph, Francis Kim, Tata Nageswara Rao, Amy Wagers, Tracy Heng, Michio Painter, Jeffrey Ericson, Scott Davis, Ayla Ergun, Michael Mingueneau, Diane Mathis, Christophe Benoist

## Abstract

Studies in mouse models have played an important role in shedding light on human hematopoietic differentiation and disease. However, substantial differences between the two species often limit the translation of findings from mouse to human. Here, we complement our previous comparative transcriptomics analysis of the human and mouse immune systems by assessing the conservation of co-expression of genes. By comparing previously defined modules of co-expressed genes in human and mouse immune cells based on compendia of genome-wide profiles, we show that the overall modular organization of the transcriptional program is indeed conserved across the two species. However, several modules of co-expressed genes in one species dissolve or split in the other species, indicating loss of co-expression. Many of the associated regulatory mechanisms – as reflected by computationally inferred *trans*-regulators or enriched *cis*-regulatory elements – are conserved across the two species. Nevertheless, the degree of conservation of regulatory mechanisms is lower than that of expression, suggesting that distinct regulation may underlie some conserved transcriptional responses.

## Introduction

The differentiation of hematopoietic stem cells into blood and immune cells in humans and other mammals is under extensive study, but less attention has been paid to the transcriptional circuitry controlling this process. Studies of human immunology typically focus on cells isolated from peripheral or cord blood or bone marrow, with in-vivo studies being limited largely to monitoring clinical manipulations, such as vaccinations or transplantations. In contrast, studies on mice, including those on immune-deficient mice transplanted with human hematopoietic cells, are usually performed in vivo [1]. Nonetheless, since all major immune cell populations are shared by human and mouse, the mouse is regarded as an important model organism for human immunology.

In the above context, many studies have suggested caution when translating findings from mouse to human [2–5], since there are substantial differences between the two species*—Mus musculus* and *Homo sapiens*. Biologically, these differences include such important factors as life span, total number of cell divisions [6], differences in a number immune cell markers [7], different physiological phenotypes caused by deficiencies in the ‘same’ (orthologous) gene in the two species (e.g., MYD88 [8], STAT5B [9]), and the presence of certain genes in only one of the two species, usually due to species- or clade-specific expansion or contraction of multi-gene families [7]. Some additional problems further complicate the translation of findings from mouse to human, as follows: First, significant disparities arise from differences in experimental protocols in that mouse studies – in contrast to those in natural human populations – are conducted primarily in inbred mouse strains and are controlled for age, environmental exposure and other confounding factors. Second, some mouse models of human disease and therapy are not readily applicable for clinical applications (e.g., asthma [10]). Third, in some cases, there may be differences during differentiation in the expression of the markers used to define the ‘same’ cell populations in the two species (e.g., in hematopoietic stem cells [3]). And, finally, analyses of mouse cells span the range of lymphoid organs, whereas studies of human material are usually limited to blood.

Despite the above challenges, it is now evident that knowledge of transcriptional profiles may indeed be exploited in the translation of findings from mouse to human, since studies in a variety of species and cell types have shown that transcriptional profiles may be used to provide a detailed view of the molecular and functional states of cells, to help identify relevant regulatory mechanisms [11, 12], and to pinpoint important differences between species [13–16]. In particular, comparing the modular organization of expression profiles and regulatory programs may provide important insights into both the conservation and the divergence of immune systems: Whereas the classical orthologous transcription factors are known to play conserved key roles in the differentiation of both human and mouse immune systems (e.g., RUNX1 and TAL1 in hematopoietic stem cells, PAX5 and EBF1 in B-cells, and TCF1 and GATA3 in T-cells), recent studies on other cell types have suggested that as many as 50% of the interactions between proteins coding for transcription factors may not be conserved between human and mouse [17] and that there are marked differences in the actual binding of transcription factors to DNA between human and mouse [18] and between other vertebrate species [19]. In particular, it has been estimated that 32%–40% of human functional regulatory sites are not functional in rodents [20].

In recent years, the way to systematically decipher the regulatory circuitry controlling hematopoiesis has been opened by the generation of two compendia of gene expression in a range of multipotent and differentiated cell types across human [21] and mouse [22] immune lineages. We previously used these two datasets to show: that human and mouse transcriptional profiles of orthologous lineages are globally similar, that signatures of lineage-specific gene expression are shared, and that expression patterns of most genes are conserved [23]. In that study, we pinpointed genes with different expression patterns in human and mouse that had not previously been reported and validated some of them experimentally [23]. However, we did not map modules of co-expressed genes between human and mouse. This mapping of modules, which is a natural extension of our previous analysis, teaches us about the similarities and differences in immune differentiation in human and mouse, particularly with regard to processes that are different or differently regulated, as discussed below.

Here, we compared the modules of co-expressed genes, which had been defined previously using the data of the human [21] and mouse [22] compendia, and showed that there is a significant similarity in the modular organization of the transcriptional program in human and mouse and a correspondingly significant overlap in the underlying regulatory programs, as defined by the inferred active regulators and associated *cis*-regulatory elements in the promoters of target genes.

## Results

### Availability of transcriptional maps of the human and mouse immune systems

We compared a compendium of the human immune system, known as the differentiation map (D-map) [21], with the mouse Immunological Genome (ImmGen) compendium [22]. D-map [21] consists of 211 samples obtained from 38 cell types (4-8 samples per cell type), measured on Affymetrix U133A arrays (22,268 probesets). The ImmGen compendium[22] consists of 802 samples obtained from 244 immune cell types (~3 samples per cell type), measured on Affymetrix MoGene 1.0 ST arrays (25,194 probesets, excluding control and unassigned features). We mapped 10,248 one-to-one orthologs between the two systems (**Materials and Methods**). Importantly, despite the differences in the design of the two studies, gene expression across the common lineages [hematopoietic stem and progenitor cell (HSPC), granulocytes (GN), monocytes (MO), dendritic cells (DCs), B-cells, natural killer (NK) cells, and T-cells] was shown to be similar [23].

### Co-expression in most modules, especially those highly expressed in stem and progenitor cells, is significantly conserved

We compared the overall organization of the transcriptional programs in human and mouse against a background of previous studies that indicated a modular organization for transcriptional programs in yeast [24] and humans [11], including the human hematopoiesis compendium [21]. Such studies also showed that in some cases modules of co-expressed genes are conserved across species, even kingdoms [25, 26]. In principle, modules can be conserved at several levels, including (**1**) conservation of gene membership (orthologous genes assigned to the ‘same’ co-expression module); (**2**) conservation of both membership and expression pattern (orthologous genes with conserved expression profiles under comparable conditions); or (**3**) conservation of a part (‘core’) of a module across two species (membership and/or expression), with additional genes in the module in one of the species, but not in the other.

Focusing first on conservation of co-expression (**1**, above), we examined modules of co-expressed genes defined independently for each compendium in its entirety (**Materials and Methods** [21, 27]). Since noise in modules reconstructed independently may somewhat reduce (but not increase) the degree of observed conservation, we assessed, for each module defined in one of the two species (i.e., human or mouse), the degree of co-expression of its members’ orthologs in the other species (i.e., mouse or human, respectively), independently of the module assignment in that other species (i.e., mouse or human, respectively) (**Fig. 1**). For this purpose, we used a Z_summary_ score (**Materials and Methods**) of module preservation, previously suggested for use across datasets and species [28]. The Z_summary_ score combines several measures of module conservation in two different datasets to reflect, for each module, the degree of co-expression of its members’ orthologs in the other species and their distinction from genes not in the module. A Z_summary_ score > 2 indicates significant conservation. The Z_summary_ statistic estimates conservation of co-expression more sensitively than a simple overlap measure between modules (below), which can be biased by the hard partitioning of similarly expressed genes to different modules with somewhat similar expression patterns. By applying this methodology, we found that 41 of the 80 human modules tested are conserved (2 < Z_summary_) in mouse and 23 of the 67 mouse modules tested are conserved in human (**Fig. 1b and c, Supplementary Table 1**). In most cases, both the co-expression and the actual expression pattern of the module’s genes are conserved, as reflected by the modules’ conservation of expression (COE) scores, calculated, according to [23], as the Pearson correlation coefficient between the mean module expression in the common lineages in both species (mean COE > 0.5 in 33 of 41 human conserved modules and 19 of 23 mouse conserved modules; option **2** above). The most highly conserved human module (module 823, Z_summary_ > 10) consists of genes whose expression is down-regulated with differentiation, suggesting particular conservation of the stem and progenitor transcriptional program. In eight human modules and four mouse modules co-expression is conserved, but the actual expression pattern is not (COE < 0.5). Thus, notably, module conservation does not necessarily imply conservation of the underlying regulatory program. Indeed, there is substantial evidence in microorganisms [29, 30] that co-expressed modules can be conserved across species, even when their underlying regulatory mechanisms diverge substantially.

**Figure 1.**
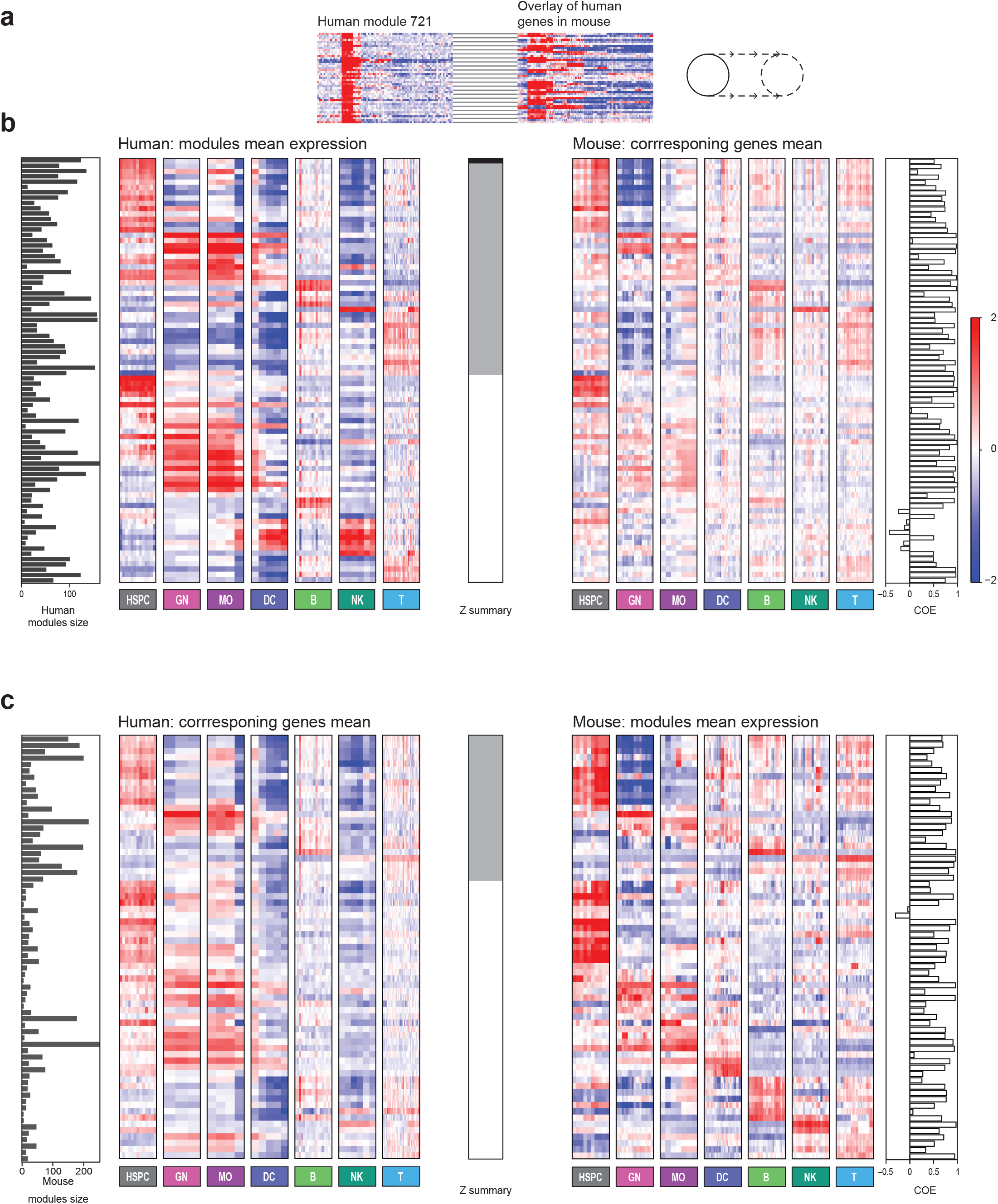
Conservation of co-expression in human and mouse immune system differentiation transcriptional programs. (**a**) Expression of each gene in human module 721 (left), as an example, compared to that of its ortholog in the mouse (right). (**b**) Mean expression profiles of all human modules (left heatmap) and of their matching sets of one-to-one orthologs in mouse (right heatmap). The bar chart on the left shows the human module size; the grayscale bar in the middle of the Figure is the Z_summary_ score reflecting module conservation (black: Z_summary_ >10 highly conserved; gray: 2<Z_summary_<10 conserved; white: Z_summary_<2 not conserved); the bar chart on the right is the COE between the mean expression of the human module and the mean expression of the mouse orthologs of its members. (**c**) Mean expression profiles of all mouse modules (right) and their matching sets of one-to-one orthologs in human (left). The bar chart on left shows the mouse module size; the grayscale bar in the middle is the Z_summary_ score of the module conservation (gray: 2<Z_summary_<10 conserved; white: Z_summary_<2 not conserved); the bar chart on the right is the COE between the mean expression of the module and the mean expression of the human orthologs of its members. (c) is analogous to (b) but projects from mouse modules to human genes (c) rather than from human modules to mouse genes (in b).

### Modular organization of the immune transcriptional program is conserved

While individual genes may be members of a module in one species but not in the other, the overall modular organization of the program may be maintained, as reflected by 61 pairs of human and mouse modules with significant overlap [**Fig. 2a and d**, hyper-geometric test, Benjamini Hochberg false discovery rate (FDR) 10%]. Those 61 pairs of human-mouse modules include 46 distinct human modules (**Fig. 2b and c**, left) and 31 mouse modules (**Fig. 2b and c**, near right), with overlaps ranging from 14% to 80% (**Fig. 2b**, far right bar chart).^1^ In most of the significantly overlapping module pairs, the expression patterns of the two modules are highly similar (median COE of module means 0.69; **Fig. 2b**, far right bar chart, **Supplementary Table 2**). However, there are cases in which only the co-expression is conserved, e.g., human module 673, induced in B-cells, and mouse module 32, induced in dendritic cells, which overlap significantly (**Fig. 2c**, bold/asterisks).

**Figure 2.**
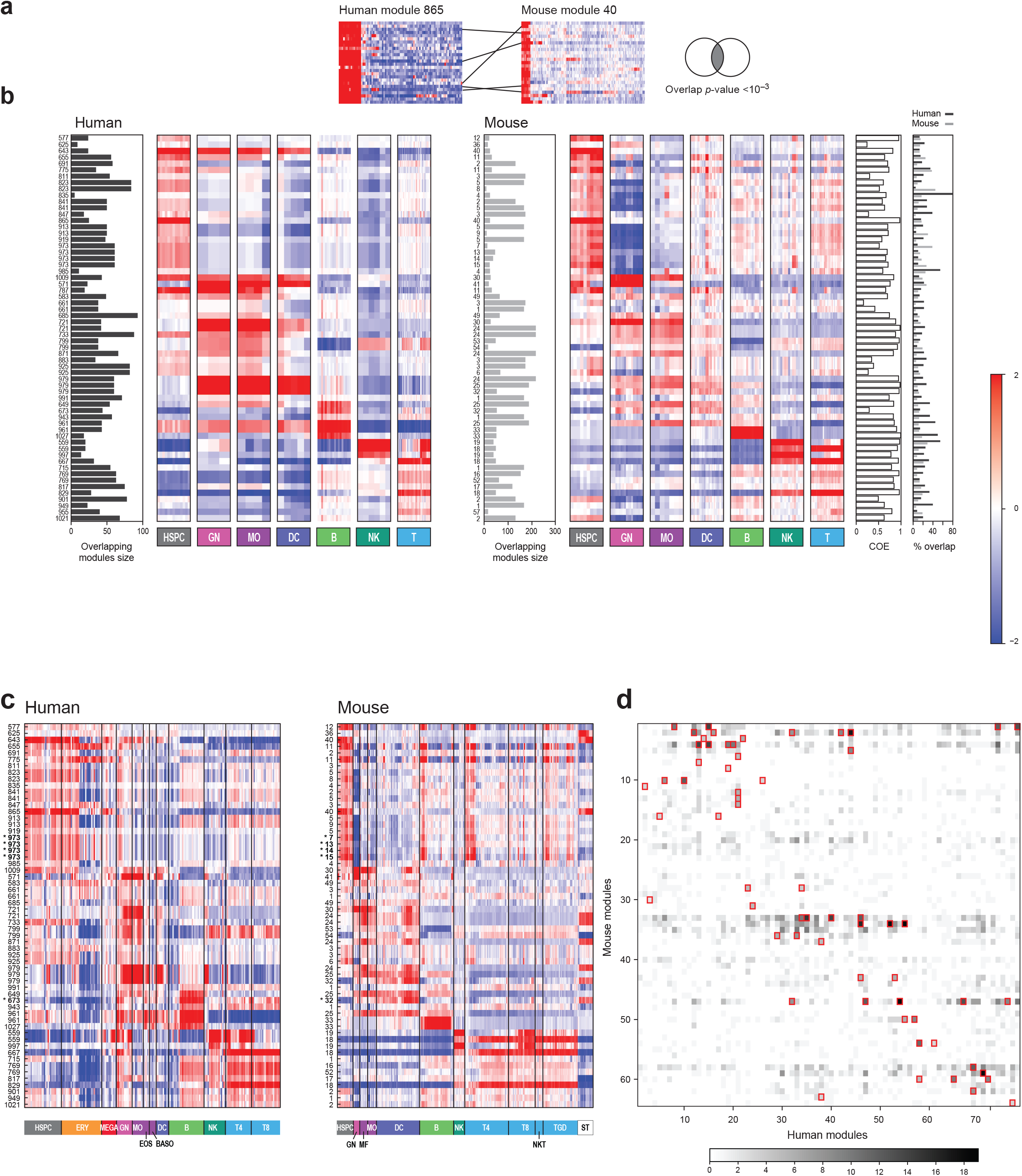
Conservation of modularity in human and mouse immune system differentiation transcriptional programs. (**a**) The gene membership in each human module (e.g., human module 865, left) and mouse module (e.g., mouse module C40, right) is compared on the basis of orthology (black lines), and the significance of the degree of overlap is estimated with a hyper-geometric test (Venn diagram, far right). Heatmaps of mean expression profiles of the common lineages (**b**) or all cell types (**c**) are shown for the 61 module pairs that significantly overlap between human (left heatmap) and mouse (right heatmap). Module pairs are sorted according to human lineage with maximal expression. Bar charts to the left of each expression matrix display module size. Bar charts on the far right are the COE between the mean expression of the matching modules (white bars) and the percentage of genes in the overlap of the human (black) and mouse (gray) modules. The expression patterns in the module pairs are typically conserved, but with some exceptions (e.g., human module 673, induced in B-cells, and mouse module C32, induced in dendritic cells). Module pairs discussed in text are marked with bold and asterisks. (**d**) Number of overlapping orthologs (grayscale color bar, right) between every pair of human and mouse modules. Significantly overlapping module pairs are indicated with red rectangles.

In some cases, the lack of one-to-one correspondence between modules is a result of the differences in cell populations profiled in the two compendia, which lead to more refined patterns in one species than in the other. For example, human module 973 [21], which consists of genes induced in HSPC and early erythroid progenitors, significantly overlaps with four mouse modules (7, 13, 14, and 15), which are all induced in HSPC but differ in expression in specific progenitors and CD8 T-cells measured only in mouse (**Fig. 2c**, bold/asterisks).

### Characterization of divergent modules

The lack of conservation – as reflected by low Z_summary_ scores – for 39 (49%) human and 44 (66%) mouse modules may be attributed to different underlying factors. These include (**Fig. 3a**): (**1**) lack of any co-expression in the other species, possibly due to differences in cell types between the compendia, resulting in module dissolution; (**2**) a relatively small conserved ‘core’ accompanied by distinct genes in each species; and (**3**) separation into several expression patterns in the other species. In addition, in some cases, the underlying clustering may over- or under-split genes into modules. To test the prevalence of each of these (possibly overlapping) possibilities, we manually inspected the expression patterns of all human modules in mouse, and vice versa.

**Figure 3.**
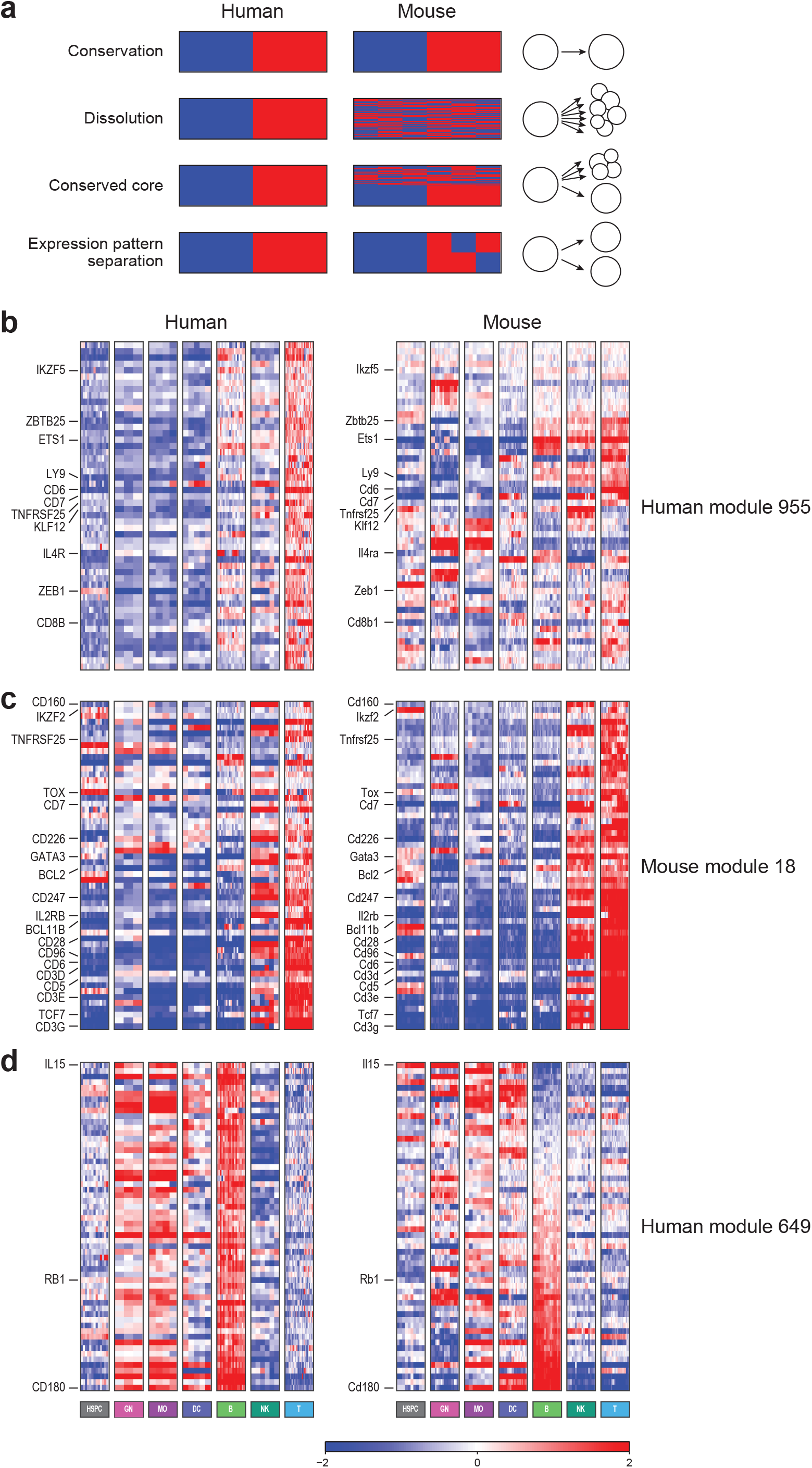
Patterns of module conservation and divergence. (**a**) Schematic visualizations of possible evolutionary patterns when comparing a module in one species (left) to the corresponding one-to-one orthologs in the other species (right). Top to bottom: conservation, dissolution, partial conservation of a core, and separation into multiple modules. In each case, schematic expression patterns in matching lineages in the two species are shown. (**b**) Human module 955 (left) is dissolved in mouse (right). Genes are sorted by their orthologs’ assignment to mouse modules. (**c**) The core of mouse module 18 (right) – induced in T-cells and also in some genes in NKs – is conserved in human (left). Genes are sorted by their mean expression in mouse T-cells. (**d**) Human module 649 (left) – induced in B-cells, granulocytes (GN) and monocytes (MO) – is split in mouse (right), with some orthologs being induced in B-cells, and others, only in myeloid cells. Genes are sorted by their mean expression in mouse B-cells. In **b-d**, genes are mean centered (color scale, bottom). DC = dendritic cells; NK = natural killer cells

The inspection revealed that four of the human non-conserved modules and fifteen of the mouse non-conserved modules are induced in cell types that are either absent or under-represented in the compendium of the other species (option **1** above). These include four modules induced specifically in human erythroid progenitors (human modules 727, 637, 889, 895), and mouse modules induced in stromal cells (modules 35-39, 44, 73, 80), T-cell progenitors (57), specific myeloid cell types (30-31, 58, 65, 68) or gamma delta T-cells (56).

Co-expression of the genes of these modules is thus cell-type specific. Additional profiling in the relevant cell types may identify similar co-expression in the corresponding species. Two non-conserved modules cannot be attributed to the lack of comparable populations, namely, human modules 859 and 955, which are induced in T-cells but whose orthologous genes in mouse display different expression patterns (**Fig. 3b, Supplementary Fig. 1a**).

In 19 human and 21 mouse modules, a ‘core’ (>50% of the module genes) is conserved (COE > 0.5) between the species (option **2** above). In many of these cases, these ‘core’ genes have higher maximal expression levels than the other genes in the same module (e.g., in mouse modules 18, 25 and 33, **Fig. 3c, Supplementary Fig. 1b**), consistent with the higher COE levels of highly expressed genes [23].

In a few cases, the orthologs of a module’s members in one species are partitioned into two distinct expression patterns (option **3** above), possibly reflecting only a portion of the pattern in the original module. For example, the genes in human modules 649 and 673 are induced in both B-cells and some myeloid cells, whereas their orthologs in mouse are induced in either myeloid or B cells but in most cases not both (**Fig. 3d, Supplementary Fig. 1c**).

### Assessing conservation of regulatory mechanisms in immune cells

The above findings lead to the question of whether the evolution of regulatory mechanisms correlates with that of the organization of the transcriptional response in the immune system. In particular, do conserved regulatory mechanisms underlie the conserved expression patterns of modules? On the one hand, studies on the evolution of gene expression in other organisms – especially yeasts – have suggested that even modules with strong conservation of expression [30] and co-expression [29, 31, 32] can be associated with distinct regulatory mechanisms in different species. In mammals, recent studies of transcription factor binding have also demonstrated substantial evolutionary turnover in transcription factor-DNA interactions, even in genes with strong functional and expression conservation [18]. On the other hand, it has been demonstrated experimentally that the functions of many known regulators of human and mouse immune system differentiation are conserved [33], suggesting that the same regulators orchestrate the process in human and mouse. In particular, turnover in the specific position of a *cis*-regulatory element or a physical binding event may exist even when the gene target or biological process controlled by a transcription factor remains conserved.

We have previously shown that the expression pattern of most genes encoding transcriptional regulators, including known master regulators, is indeed conserved, which is a prerequisite for the conservation of regulation [23]. To study the extent of conservation of regulatory mechanisms between human and mouse, we examined two aspects of the immune system regulatory program, as reflected in the transcriptional compendia and associated modules (**Fig. 4a**), namely: (**1**) inferred *trans*-regulatory associations between regulators and targets, as predicted by computational modeling of gene expression patterns; and (**2**) *cis*-regulatory elements enriched in promoters of module members or lineage-signature genes.

**Figure 4.**
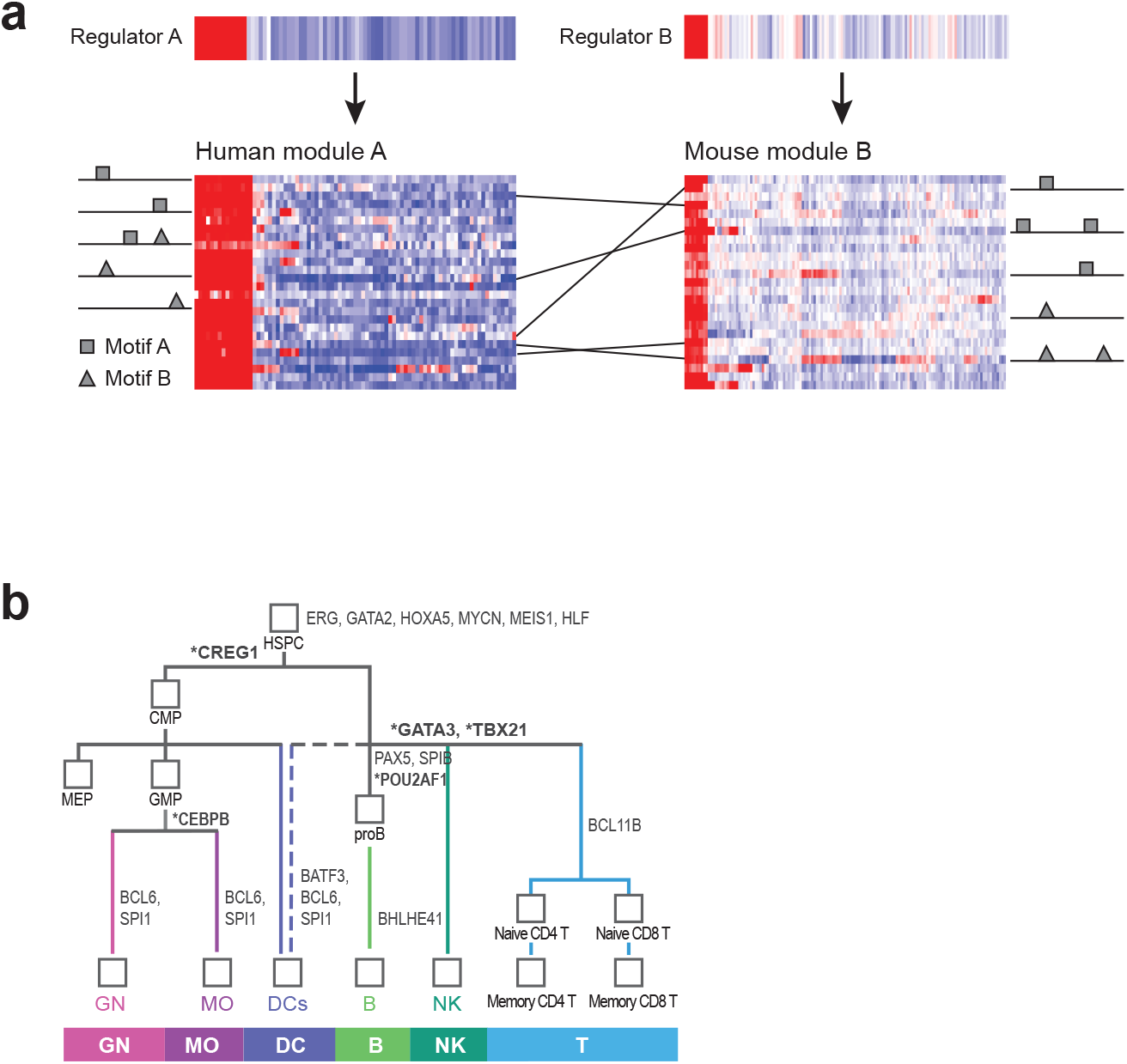
Regulation of human and mouse immune system differentiation is largely conserved. (**a**) Schematic representation of several possible levels of conservation of regulation in human and mouse: expression of regulators (top), regulators associated with modules (arrows), and *cis*-regulatory elements enriched in promoters of module genes (rectangles and triangles, right and left). (**b**) Conserved lineage association of regulators. Shown is the lineage tree with selected immune system regulators with conserved expression (COE>0.8). Conserved lineage associations according to the regulatory models are presented in bold and indicated with asterisks. The full set of conserved lineage associations is provided in **Supplementary Table 4**.

### Many of the regulators predicted by computational modeling are conserved

To identify active *trans* regulators associated with each module, we used Ontogenet, a method that combines linear regression with the tree structure of the dataset to predict the set of transcriptional regulators that would best account for each module’s expression [27]. Ontogenet has already been applied to mouse modules with 580 candidate regulators and has identified 480 regulators (7-45 per module, median 16) [27]. Here, we applied Ontogenet to the human modules independently, using 394 candidate regulators (**Supplementary Table 3**), and identified 213 regulators in human (0-30 per module, median 15).

Of the 480 regulators chosen by the Ontogenet algorithm in mouse (‘active regulators), 155 were also chosen as active regulators in the human model. The larger number of candidate regulators and active regulators in mouse reflects the higher complexity of the mouse dataset The COE scores of the 155 active regulators chosen in both models are significantly higher than those of other genes filtered in a similar manner [one sided Kolmogorov–Smirnov (KS) test, P = 2.7*10^-6^] but not of the active regulators that are selected only in one species (one sided KS-test, P = 0.12). Furthermore, the targets and lineages associated with a few of the regulators are significantly conserved between the two regulatory programs. Of the 155 pairs of orthologous regulators chosen in both species, the targets of 13 regulators (CBX5, CIITA, FUBP1, HIVEP2, LRRFIP1, NPM1, PHB2, POU2AF1, SPI1, SPIB, TCF4, TFEC and STAT6) significantly overlap between the human and mouse models (hyper-geometric test, FDR = 10%). The number of overlapping pairs (13) is greater than the number that would be expected by chance (P<10^-3^, permutation, **Materials and Methods**). Finally, many specific regulators are conserved in their association to individual lineages (e.g., **Fig. 4b**, bold/asterisks). For each lineage, testing of the overlap of activators and repressors between human and mouse (**Supplementary Table 4**) showed that T-cell activators and repressors and DC repressors significantly overlap between the human and mouse models (hyper-geometric test, FDR = 10%, **Materials and Methods**).

Most of the pairs of modules conserved between human and mouse (40 of 61) are associated with at least one pair of orthologous regulators (**Supplementary Table 2**). For example, POU2AF1 is a regulator of the B-cell modules human 961 and mouse C33; TBX21 is a regulator of the NK-cell modules human 997 and mouse C19; and GATA3 is a regulator of the T cell modules human 667 and mouse C18. The regulators that were chosen in one species but not the other have a significantly lower range of expression in the species in which they were not selected (t-test P = 2.8*10^-3^ for mouse; P = 1.2*10^-4^ for human). Indeed, many of the discordant regulators are highly expressed in the non-common lineages (e.g., the regulatory T-cell regulator Foxp3 and the stromal specific Epas1 chosen only in mouse are expressed only in cell types not measured in the human dataset).

### Some of the *cis*-regulatory elements associated with modules and signatures are conserved

There is substantial conservation in the *cis*-regulatory elements associated with the regulatory programs in the two species, suggesting conserved regulatory mechanisms. We found that 16 of the 61 pairs of modules conserved between human and mouse are associated with at least one *cis*-regulatory element that is enriched in both members of the pair (**Supplementary Table 2**). For example, human module 973 and its orthologous mouse modules 7, 14, and 15, all consist of genes down-regulated with differentiation; they also contain many cell-cycle genes and are enriched with the binding sites of different E2F factors. E2F factors are known cell cycle regulators [34], and the expression of E2F1 and E2F3 is down-regulated with differentiation.

### *Cis*-regulatory conservation is not associated with higher conservation of expression

Surprisingly, the presence of conserved *cis* elements is not associated with significantly higher conservation of expression as measured by COE. For example, there is not a greater similarity in expression in pairs of orthologous modules that are enriched for the same sequence motif in human and mouse than in pairs of modules not having matched enriched motifs (KS test, *P* = 0.85). This finding may be due either to the relative paucity of known *cis*-regulatory elements and inaccurate prediction of their targets [35, 36] or to the presence of dense regulatory circuits [21], such that compensation for loss of *cis* regulation by one factor is awarded by gain of regulation by another factor active in the same lineage.

## Discussion

While many features of the immune system are conserved between human and mouse – such that the mouse immune system can indeed be regarded as a compelling model for human immune system differentiation – there are important known differences between the two systems, including those in transcriptional profiles (reviewed in Mestas *et al*. [7]). Nonetheless, despite the knowledge that has accumulated to date, previous studies have not systematically analyzed the extent of similarity and differences in the human and mouse regulation of transcriptional programs of the immune system.

Our previously published comparison of the two extensive transcriptional compendia showed extensive conservation across the two species of the transcriptional program at several levels, namely, in terms of global profiles and of individual genes and lineage-specific gene signatures; that study also catalogued the transcriptional differences in one-to-one and one-to-many orthologs [23]. Here, we complement that study by showing conservation and divergence at the level of modules of co-expressed genes and by comparing the underlying regulatory mechanisms controlling these programs in the two species. In addition, we show that when a regulatory module in one species is partially ‘dissolved’ in another, co-expression (and hence module membership) of genes with higher expression levels tends to be more conserved, in accordance with our previous observation that the expression pattern of highly expressed genes is more conserved [23].

The most prevalent expression pattern in the two species is downregulation with differentiation, that is, genes whose expression is high in HSPC and down-regulated in all other cell types. In accordance, many of the conserved modules contain genes expressed specifically in stem and progenitor cells. The prevalence of this expression pattern and its conservation may reflect a stronger purifying selection against changes in the HSPC transcriptional program – where changes could disrupt many differentiation paths – whereas the ‘differentiated cell types program’ may be under less selective pressure.

Module conservation is often reflected by a concomitant conservation of the associated regulatory mechanisms. In particular, the expression profiles of most regulators chosen by the regulatory models are well conserved. Indeed, there is a significant overlap between lineage regulators assigned by the two models. Furthermore, many conserved modules are also associated with conserved enrichment of regulatory elements. Nevertheless, in many cases conserved modules are associated with different *cis*-regulatory elements. Such conservation of gene expression and co-expression but divergence of underlying *cis* elements has been previously described in yeast [29, 31, 37]. Some of the discrepancies between conservation of expression and lack of conservation of *cis* elements are probably related to the lower reliability of *cis*-regulatory predictions or to differences in the sampled cell types. For example, the mouse compendium does not include a counterpart to the interferon gamma-producing CD56+ human NK cells. In other cases, the preponderance of dense regulatory circuits in hematopoiesis [21] may facilitate the divergence demonstrated in the current study.

Being aware that our comprehensive comparison of human and mouse modules of co-expressed genes and their regulation is of great interest to the immunology and evolution community, we provide all the data and analysis in a separate browser on the ImmGen portal (http://rstats.immqen.orq/comparative/comparative_search.php) so as to facilitate future studies of the evolution of genes’ regulation. It is our belief that this analysis will assist researchers in identifying both the broad similarities and fine distinctions between human and mouse in the modular structure of the immune system transcriptional program.

## Acknowledgements

We thank the members of the ImmGen Consortium for discussions, the ImmGen core team at the time of analysis (M. Painter, J. Ericson, S. Davis) for help with data generation and processing, and eBioscience, Affymetrix, and Expression Analysis for support of the ImmGen Project. We thank Leslie Gaffney for help with figure preparation, Lewis L. Lanier, Roi Gazit, and Paul A. Monach for many helpful discussions, and Noa Novershtern and Benjamin L. Ebert for help with the human dataset.

## Materials and Methods

### Datasets and preprocessing

Gene expression in mouse samples was measured on an Affymetrix array MoGen, annotation version na31. The ImmGen March 2011 release was used. This release includes 802 arrays with 22,268 probesets, as previously described [27]. Gene expression in D-map human samples was measured on an Affymetrix array U133A, annotation version na31, as previously described [21]. There are 211 arrays with 22,277 probesets. We used the ImmGen RMA normalized data and the D-map normalized and batch corrected dataset as previously published [21, 27]. When more than one probeset had the same gene symbol, only the probeset with the highest mean expression was used.

### Orthologous gene mapping and filtering

We used Ensembl COMPARA release 63 to map orthologs. Mouse ENSEMBL gene IDs were matched to 15,265 probesets on MoGen, and human ENSEMBL gene IDs were matched to 10,457 probesets on U133A. In COMPARA there are 15,678 one-to-one orthologs and 539 apparent one-to-one orthologs, resulting in 16,217 one-to-one mapped genes between the species. Of these, 10,248 one-to-one ortholog pairs were measured in the two arrays—human and mouse. Finally, only the 5,841 genes with an expression level above 120 (recommended ImmGen threshold for expression) in at least three arrays of the common samples (see [23] for details on mapping human-mouse samples, particularly **Supplementary Table 1** in that reference) in both species were included in the filtered set of one-to-one orthologs. All further analysis was performed on this set.

### Conservation of expression (COE)

The COE is a measure of agreement of expression in comparable groups of samples, i.e., lineages, between two species. We previously defined it at the gene level as follows: for each species, we first computed the median of the gene’s expression in each of the seven common lineages, and then calculated the COE of the gene as the Pearson correlation between these lineages [23]. Similarly, the COE of a module’s expression is the COE of the mean expression of the genes in the module (since variation from the module’s mean is expected to reflect noise).

### Module conservation and comparison

We used the 80 human modules as previously published [21] and 81 mouse modules from the ImmGen Consortium analysis [27]. Of these, we analyzed the 80 human modules and 67 mouse modules containing 5 or more one-to-one orthologs of the filtered set. Conservation of modules was estimated by the Z_summary_ statistic suggested by Langfelder *et al*. [28]; for this purpose, the WGCNA R package was used. The Z_summary_ statistic estimates the conservation of a reference dataset module in a test set. When testing for conservation of human modules, the full human dataset was used as the reference set, and the full mouse dataset was used as the test set (**Supplementary Table 1**). Similarly, when testing for conservation of mouse modules, the full mouse dataset was used as the reference set, and the full human dataset was used as the test set (**Supplementary Table 1**).

We calculated the overlap between each pair of human and mouse modules, and assigned a p-value for module overlap using a hyper-geometric distribution for two samples, with the genes in the filtered set that were assigned to modules in both species serving as the background.

### Comparison of model regulators

We used the mouse regulatory model defined for the mouse modules by Ontogenet [27] and created a distinct regulatory model for human by applying Ontogenet [27] independently to the human modules. Since the original publication on the human dataset [21] used a different method to create a regulatory model, we avoided a direct comparison to it so as to prevent artifacts due to the difference in the analysis tools used in the two species.

Ontogenet was specifically devised to address some of the challenges – and leverage some of the unique power – of studying transcriptional programs in cell lineages. Thus, as elaborated in [27], Ontogenet has several major advantages over the method originally used in the D-map study (Module Networks [11]). First, Ontogenet can identify a whole set of ‘equivalent’ regulators, whereas the Module Networks approach would have had to choose (somewhat arbitrarily) only one representative. Allowing multiple regulators is more consistent with the dense interconnected nature of regulatory circuits that control cell states. Second, Ontogenet allows us to choose a regulator in a context-specific manner, assuming that it may be relevant to the regulation of a gene module only in some cells in the lineage, but not others. Third, Ontogenet uses the lineage tree to guide its search for a regulatory program, by preferring (but not mandating) models in which ‘close’ cells in the lineage share regulatory mechanisms.

The candidate regulators for the mouse model (from which the method chooses active regulators) followed those previously reported for human [21], with minor adjustments, as described in [27]. We used their one-to-one orthologs as candidate regulators for the human model (**Supplementary Table 3**). The targets of a regulator were defined as the union of all modules to which it was assigned. We tested for significant overlaps between targets of the 155 regulators chosen by Ontogenet in the two species by using a hyper-geometric test with an FDR of 10%. The background comprised all genes assigned to modules in both species and included in the filtered one-to-one orthologs set. To test whether the number of orthologous regulator pairs whose targets significantly overlap (13-out-of-155) is higher than would be expected by chance, we permuted the regulatory interactions of the 155 common regulators in one species while preserving the regulatory interactions at the module level and repeated the calculation 1,000 times. When comparing regulators of significantly overlapping (orthologous) modules, we used a hyper-geometric test to test the significance of the overlap with an FDR of 10%, using as the background all 155 candidate regulators that are included in the filtered one-to-one orthologs set and had been chosen by Ontogenet.

Ontogenet also provides for each lineage a list of activators and repressors, based on their average regulatory weights across all cell types in the lineage and all modules. For each lineage, we used a hyper-geometric test with an FDR of 10% to test the significance of the overlap of the lineage activators in human and mouse and, similarly, of the lineage repressors in the two species. The background comprised all regulators that had been chosen by the model in either species and are included in the filtered one-to-one orthologs set. FDR was applied to the p-values of all 14 regulators groups (7 lineages, activators and repressors for each).

### Motif enrichment in modules

Motif scanning and motif scoring threshold were performed as previously described [27], resulting in a MAX-LOD(*i,k*) score for each motif *k* in each gene *i*. This score reflects the best motif instance over the entire promoter region. For each module of genes *M* and each motif *k*, we computed the p-value for enrichment, *p_e_*(*M,k*), of the motif in the module, compared to the entire set of genes assigned to modules in that species serving as the background. An enrichment of a motif in a module results in higher than expected *MAX-LOD* scores for the genes in that module: To capture this effect, we computed the p-value by comparing the *MAX-LOD(i,k)* scores for all genes *i* in the module *M* and the scores for the entire set of genes assigned to modules in that species by performing a one-sided rank-sum test. We then employed an FDR of 5% on the entire matrix of p-values *p_e_*(*M,k*) and declared as significant hits all pairs of modules and motifs that were assigned p-values lower than the FDR threshold.

### Estimating the effect of conserved *cis*-regulatory elements on expression

We used a one sided KS-test to estimate whether the 286 human and mouse module pairs whose genes are enriched for the same motif are more similar in expression pattern, as defined by a COE measure, than would be expected by random. The background comprised a COE distribution generated from 1,000 repeats of the COE of 286 randomly selected modules from all human modules enriched for any motif and 286 randomly selected modules from all mouse modules enriched for any motif.

### Multiple comparison control

The Benjamini Hochberg false discovery rate (FDR [38]) procedure was used to control the false discovery rate at 5% or 10%, as stated.

---

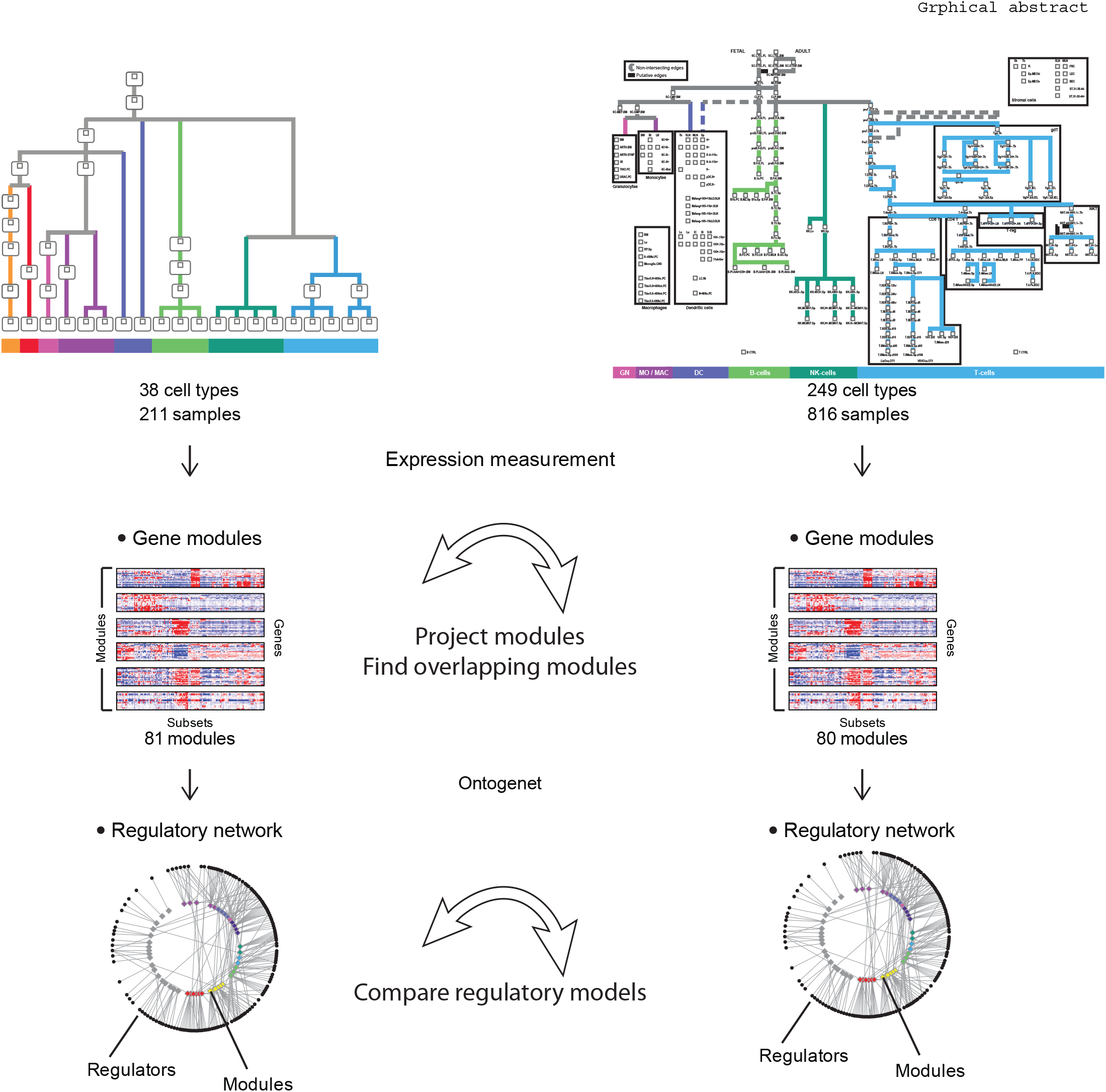

**Supp. Figure 1.**
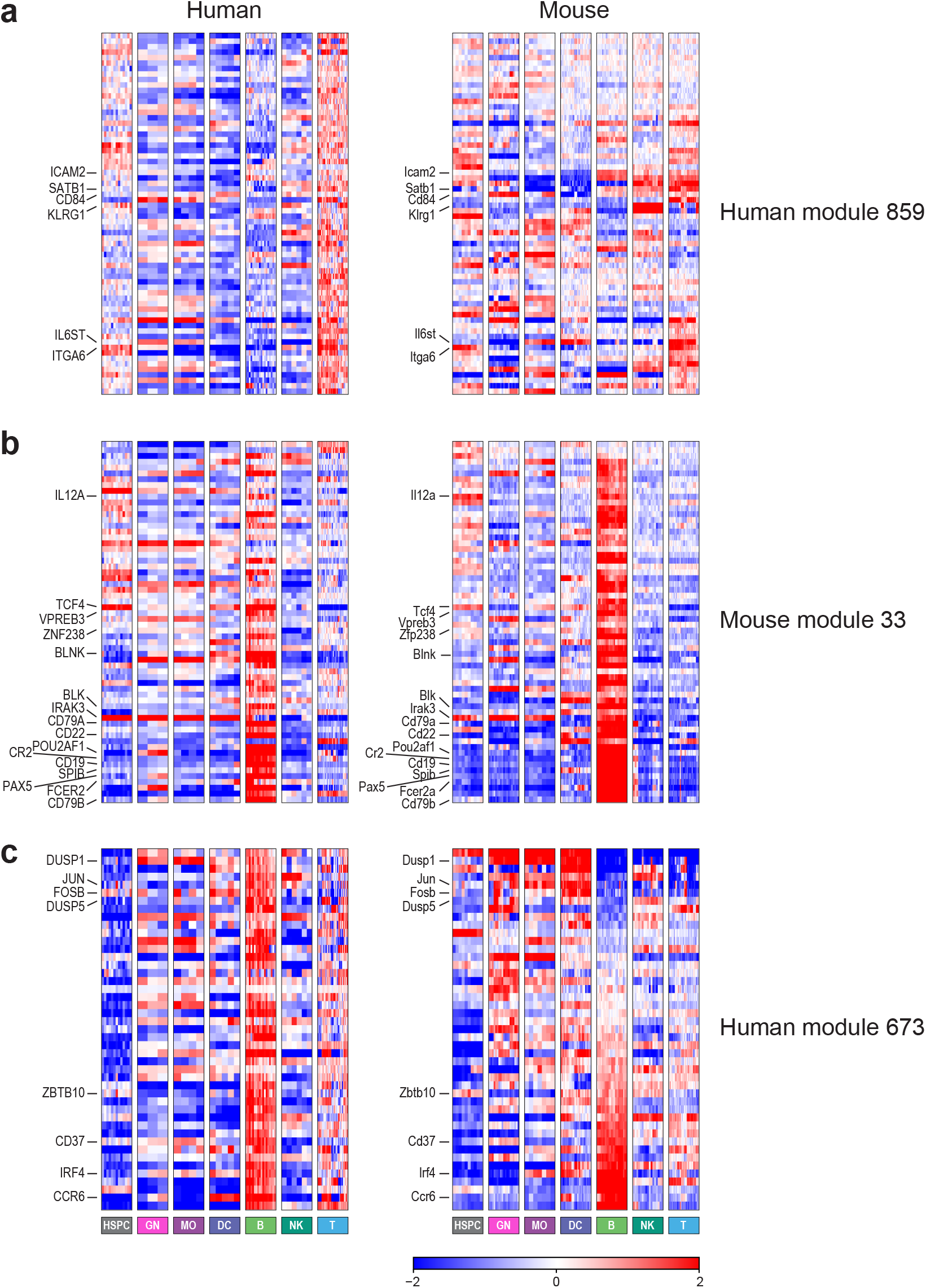

## Footnotes

1 Note that one human module may significantly overlap more than one mouse module, and vice versa.

